# Somatic variant analysis of linked-reads sequencing data with Lancet

**DOI:** 10.1101/2020.07.04.158063

**Authors:** Rajeeva Musunuri, Kanika Arora, André Corvelo, Minita Shah, Jennifer Shelton, Michael C. Zody, Giuseppe Narzisi

## Abstract

**Summary:** We present a new version of the popular somatic variant caller, Lancet, that supports the analysis of *linked-reads* sequencing data. By seamlessly integrating barcodes and haplotype read assignments within the colored De Bruijn graph local-assembly framework, Lancet computes a barcode-aware coverage and identifies variants that disagree with the local haplotype structure.

**Availability and Implementation:** Lancet is implemented in C++ and is available for academic and non-commercial research purposes as an open-source package at https://github.com/nygenome/lancet.

## 1 Introduction

Local assembly approaches have become the preferred solution to increase variant calling power compared to read alignment-based methods (Narzisi *et al*., 2014; Cooke *et al*., 2018; Chen *et al*., 2016; Wala *et al*., 2018; Rimmer *et al*., 2014; Mose *et al*., 2014; Li *et al*., 2013; Narzisi et al., 2014; Cooke et al., 2018; Chen et al., 2016; Wala et al., 2018; Rimmer et al., 2014; Mose et al., 2014; Li et al., 2013; Benjamin et al., 2019). Lancet is a somatic variant caller that leverages local assembly and joint analysis of tumor-normal paired data using region-focused colored de Bruijn graphs, with on-the-fly repeat composition analysis and a self-tuning *k*-mer strategy. This results in: reduced reference bias; improved ability to detect variations that significantly diverge from the reference chromosome representations; reduction in the scale of the analysis, leading to increased power and sensitivity to detect variants through localized, comprehensive graph exploration; and dynamic adjustment of calling behavior according to the sequence conditions of each genomic region. We previously demonstrated the superior accuracy of this approach, especially for InDels, compared to widely used somatic variant callers (Narzisi *et al*., 2018). Moreover, a recently published and independent testing (Köster *et al*., 2020), showed Lancet to outperform state-of-the-art methods particularly in the detection of twilight zone InDels (30-250 bp).

Recently developed “linked-reads” technologies (Wang *et al*., 2019; Zheng *et al*., 2016) are capable of extending the Illumina system to include long-range information at low cost and with minimal sample requirements, using barcodes to identify reads sampled from the same molecule. Yet, for this new data type to achieve its full potential and benefit the larger cancer research community, new computational tools that can leverage it must be developed. High quality germline variant analyses have been shown to be achievable with 10xGenomics (https://www.10xgenomics.com/) linked-reads, with the added value of retaining long-range information for phasing and structural variant calling resolution (Marks *et al*., 2019). However, as we show here, off-the-shelf somatic variant callers perform poorly on these data, due to presence of artifacts and sources of error, including high rate of chimeric reads and discordant read pairs, affecting the detection of low frequency variants. Moreover, the phasing algorithm implemented in the LongRanger suite from 10xGenomics assumes a diploid model, hence lacking direct support for the clonal structure of cancer genomes. Specialized algorithmic solutions are therefore needed to handle these new error models, as well as phasing information, to provide accurate somatic variant detection. Here, we describe how we extended Lancet to support the analysis of linked-reads sequencing data and compare its performance to established somatic variant callers today.

## 2 Methods

Barcodes from the linked-reads can be used to confidently infer which variants are present on the same molecule, and therefore phased within the same chromosome copy, or subclone in cancer genomes (**Supplementary Fig. S1**). If a real mosaic variant arises near a heterozygous mutation, it should always be found in conjunction with only one of the two alleles and should never appear on reads with the other allele. This generates three haplotypes in bulk sequencing supported by the barcodes configuration (**Supplementary Fig. S2**). Consequently, genuine somatic calls should never manifest as mutations supported by reads on both haplotypes.

We have extended Lancet to seamlessly integrate barcodes and haplotype read assignments within the colored de Bruijn graph local-assembly framework. Specifically, each node (*k*-mer) is augmented to store the set of barcodes and haplotype groups associated with the linked-reads from where each *k*-mer was extracted (**Supplementary Section S1**). Then, Lancet uses this information to identify variants that disagree with the local haplotype structure (“haplotype filter”). Barcode-aware coverage is also computed by threading through the graph to count the number of distinct barcodes associated to each *k*-mers in the path. Similar to flagging overlapping read pairs from the same fragment, this procedure is necessary to avoid false-positive (FP) low frequency errors due to overcounting coverage of reads sharing the same barcode (**Supplementary Fig. S3**). The output VCF lists allele-specific barcodes for each called variants, allowing downstream identification of somatic mutations co-occurring on the same haplotype and/or subclone.

## 3 Results and discussion

To accurately evaluate the performance of our approach, we created a virtual tumor-normal pair using high-coverage whole-genome linked-reads data for the NA12891 and NA12892 HapMap samples with spiked-in mutations at known sites and frequencies (**Supplementary Section S2**). By precisely knowing the true set of variants, we used the virtual tumor to estimate FP rate, inspect erroneous calls, and develop specialized filters. We first noticed several artifacts specific to the linked-reads that reduce the overall quality of the 10xGenomics data compared to standard Illumina data (**Supplementary Fig. S4**), including high rate of incorrectly marked PCR duplicates (**Supplementary Fig. S5**). Running Picard Tools MarkDuplicates on the LongRanger BAM file reduced the number of these errors by 44% and 23% for SNVs and InDels respectively (**Supplementary Fig. S6**). Lancet shows better performance and substantially lower number of FPs compared to MuTect2 and Strelka2, indicating that somatic callers developed for standard Illumina data do not handle well the different error profile of linked-reads data (**Fig. 1** and **Supplementary Fig. S7**). Distribution of the variant allele fractions (VAF) of the somatic SNVs shows that the majority of the FPs occur at low VAFs (**Supplementary Fig. S8**), limiting the detection of real low frequency somatic events.

**Figure 1.**
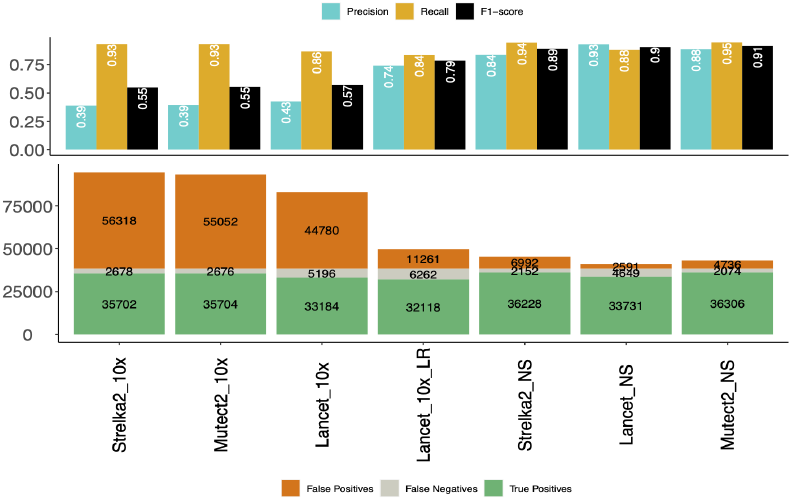
Linked-reads virtual tumors variant calling evaluation.

We then analyzed publicly available linked-reads data for the two well characterized COLO829 and HCC1954 cancer cell lines (**Supplementary Table T1**), using high confidence calls from the NYGC cancer pipeline (Arora *et al*., 2019) as the truth set. The linked-reads aware Lancet has the best balance between precision and recall, and, compared to the original Lancet on COLO829, preferentially removes 69% and 77% of the false-positive SNVs and InDels respectively, with only marginally reduced sensitivity (**Supplementary Tables T2 and T3**). Counts of supporting reads per haplotype confirm the capability of the haplotype filter to accurately classify FP events (**Supplementary Fig. S9**). Categorization of FP calls by sequence context shows enrichment for homopolymers and STRs in linked-reads data compared to standard Illumina sequencing, likely due to PCR artifacts (**Supplementary Fig. S10**).

In summary, we have enhanced Lancet to take advantage of the information available in linked-reads sequencing. Compared to current state-of-the-art somatic callers (Mutect2 and Strelka2), Lancet shows higher accuracy, currently making it the tool of choice to analyze cancer datasets sequenced with linked-reads. However, we report higher error rates for the 10xGenomics linked-reads compared to standard Illumina data, which limits the detection of low frequency mutations, especially InDels. Adjusting for molecule coverage and PCR artifacts mitigates some of the problems, but sensitivity and precision with linked-reads still remain inferior compared to employing regular Illumina data.

## Acknowledgements

We thank the Computational Biology team at the New York Genome Center for their support and suggestions and the NYGC Research Computing group for their help with data processing.

