## Supplementary Material for "Somatic variant analysis of linked-reads sequencing data with Lancet"

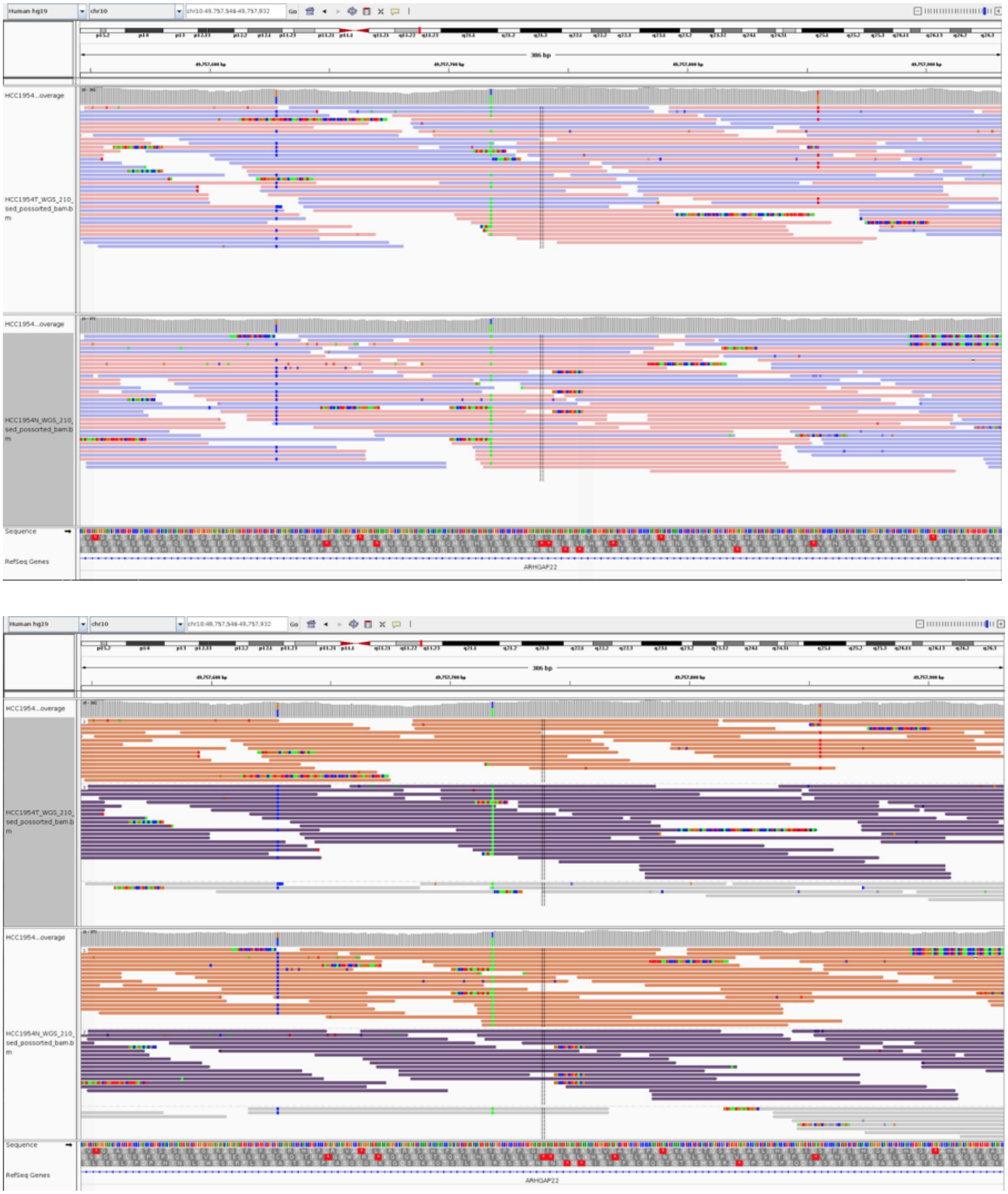

**Figure S1. Example of phasing information on a combination of germline and somatic variants.** Read alignments for two heterozygous germline variants (blue and green) and a somatic mutation (red) for a tumor-normal pair. Without phasing information (top figure) it is not possible to disentangle the relative linkage of the variants. After phasing with linked-reads (bottom figure) it becomes clear that the somatic variant does not co-occur with the nearby germline variants.

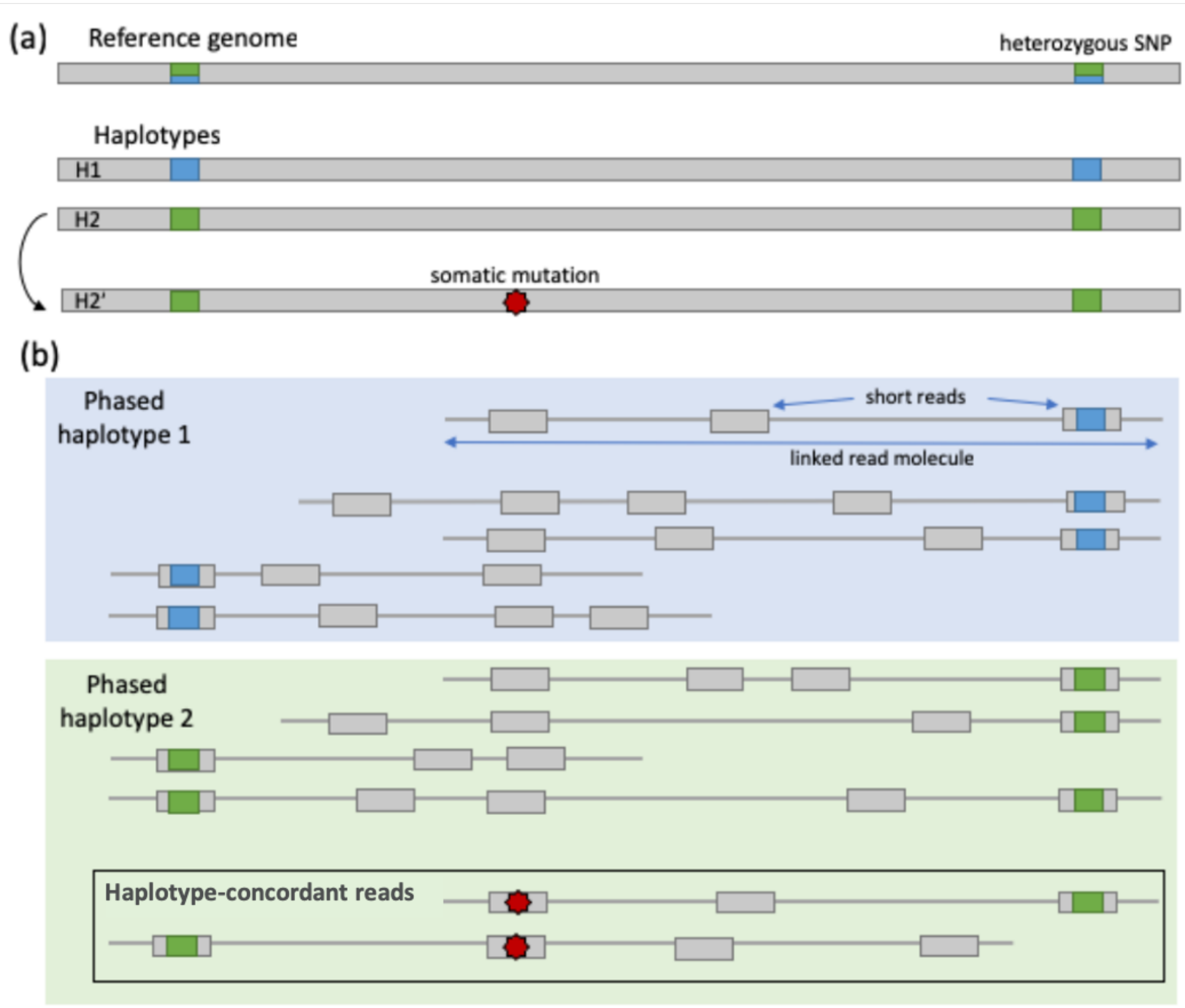


**Figure S2. Linked-reads to distinguish real somatic mutations from sequencing errors (Haplotype filter).** If a real mosaic variant (red star) arises near a heterozygous mutation it should always be found in conjunction with only one of the two alleles (green) and should never appear on reads with the other allele (blue). This generates three haplotypes in bulk sequencing. So, genuine somatic calls should never manifest themselves as mutations supported by reads on both haplotypes (e.g., phased with mutations occurring on different haplotypes). Figure from Darby et al. "Samovar: Single-Sample Mosaic Single-Nucleotide Variant Calling with Linked Reads" iScience 18, 1–10 (doi: https://doi.org/10.1016/j.isci.2019.05.037), originally adapted from Figure 3 of Dou Y, Gold HD, Luquette LJ, Park PJ. "Detecting Somatic Mutations in Normal Cells" Trends Genet. 2018;34(7):545–557 (doi:https://doi.org/10.1016/j.tig.2018.04.003).

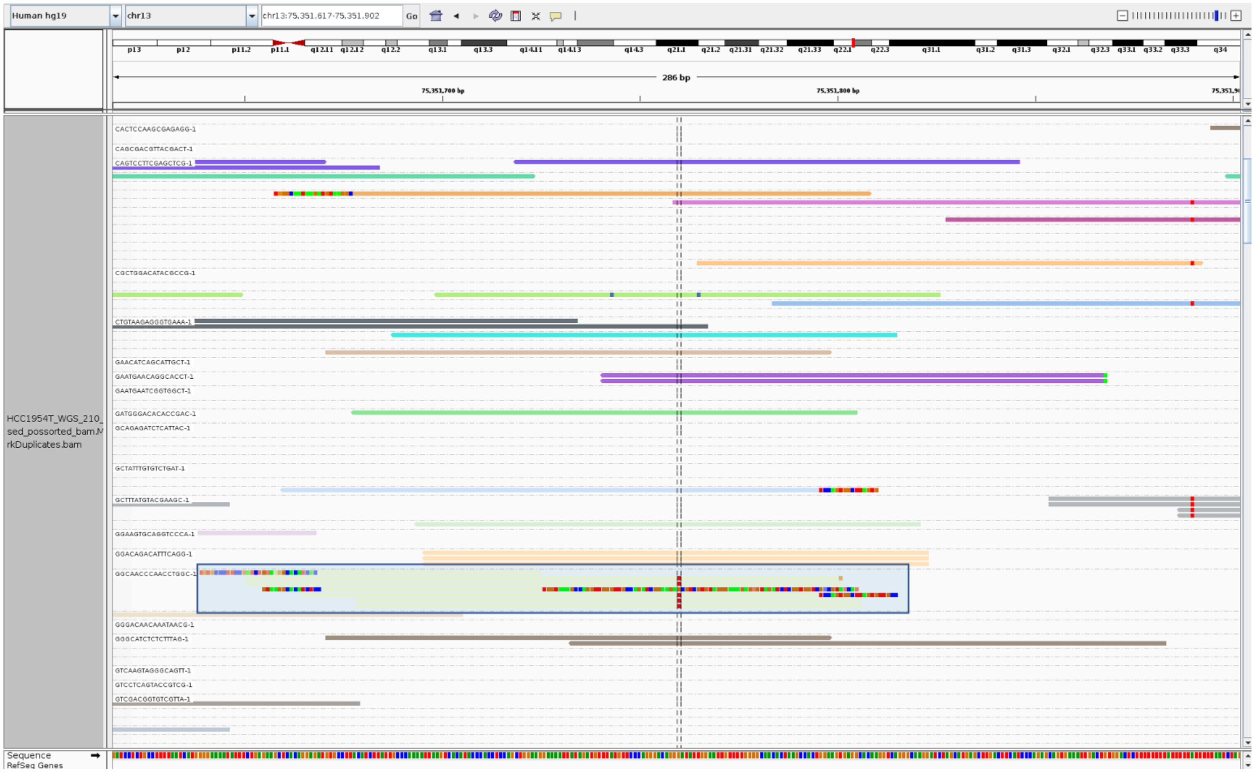

**Figure S3. 10x Genomics Linked-reads barcode artifacts.** Example of overlapping reads sharing the same barcode from one single molecule, showing the importance to avoid overcounting the number of supporting reads when computing coverage.

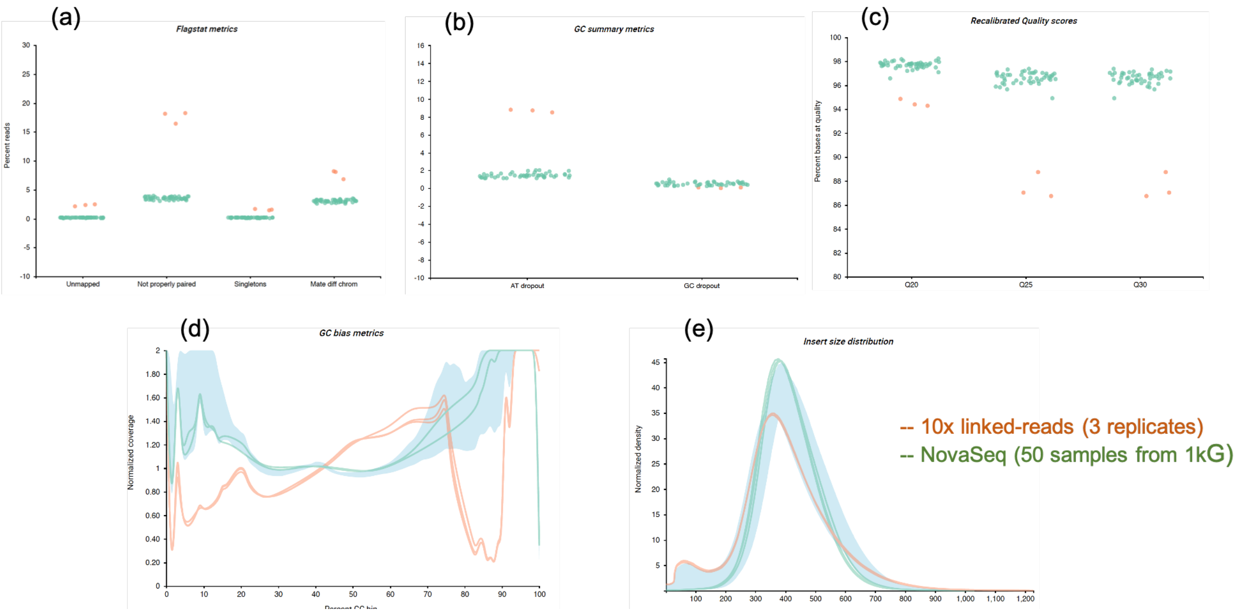


**Figure S4. 10x Genomics Linked-reads QC metrics.** (a) Read pairs flagstat metrics, (b) GC summary metrics, (c) Recalibrated quality scores, (d) GC bias, and (e) insert size distribution comparison between three 10x Genomics replicates and 50 NovaSeq samples from the 1000 Genomes project sequenced using the NovaSeq platform.

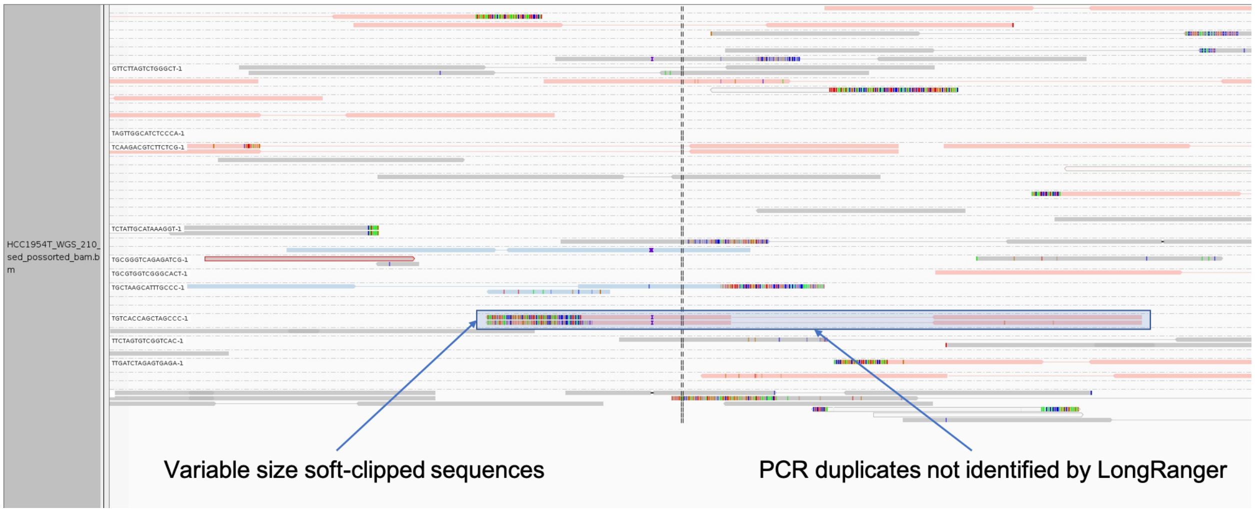


**Figure S5. 10x Genomics Linked-reads PCR artifacts.** Example of PCR duplicates not correctly identified and marked by LongRanger, possibly due to the variable size soft-clipping. Without correction this can introduce false-positive variant calls.

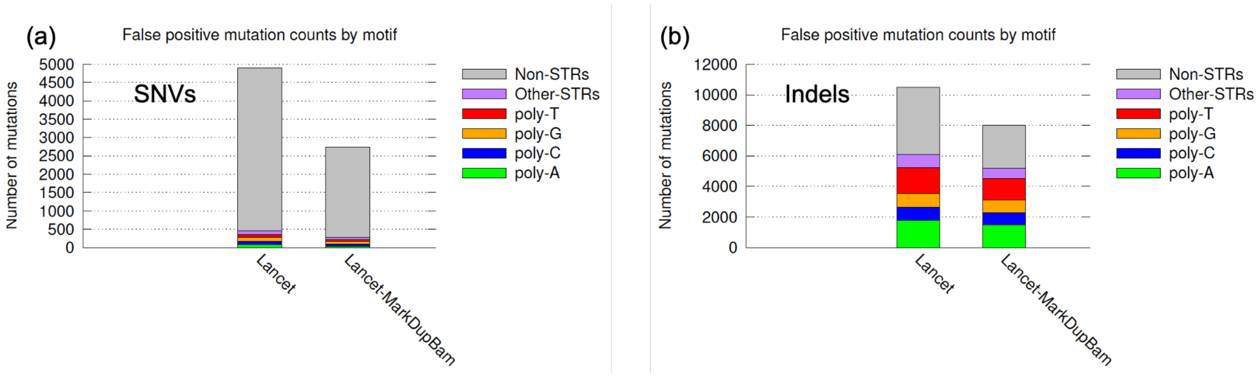


**Figure S6. Impact of PCR-duplicate correction in variant calling.** Impact of PCR-duplicate correction on the number of false positives variants called by Lancet on the linked-reads virtual tumors. Running Picard Tools MarkDuplicates on the LongRanger BAM file correctly flags read-pairs and reduces the number of false-positive calls by ~44% and ~23% for (a) SNVs and (b) InDels respectively.

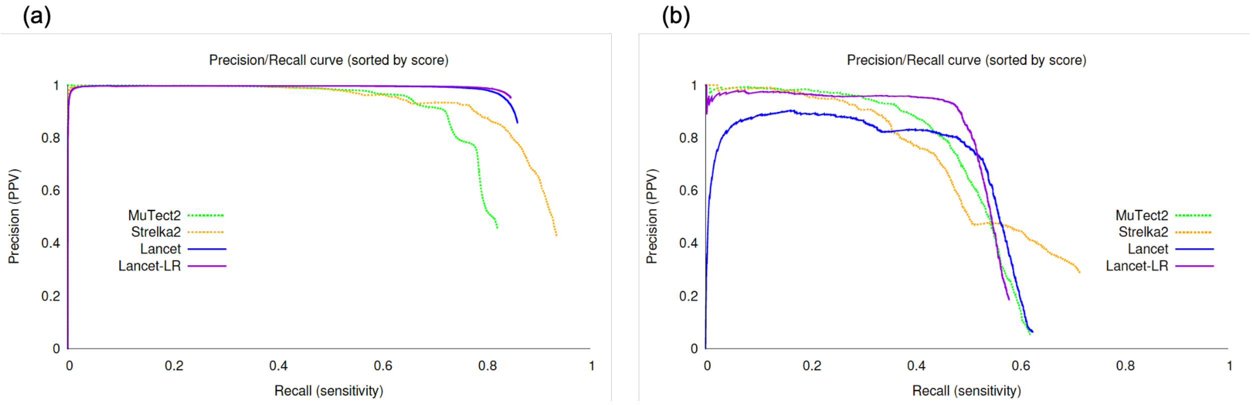


**Figure S7. Precision/recall performance on the linked-reads virtual tumor.** Precision/recall curve analysis of somatic (a) SNVs and (b) InDels called by Lancet-LR, Lancet, MuTect2, Strelka2 on the linked-reads virtual tumor. Curves are generated by sorting the mutations based on the confidence score assigned by each tool (from highest quality to lowest). Each point of the curve corresponds to the precision and recall for all the variants with confidence scores greater than or equal to a specific quality threshold. The curve for an ideal tool (no errors) would start from the top left corner and produce a straight horizontal line (with precision=1). Any deviation from a straight line is due to errors introduced by the variant calling process. Specifically, deviations at low recall rates are indicative of low performance of the scoring system adopted by the tool (false positive variants with high score).

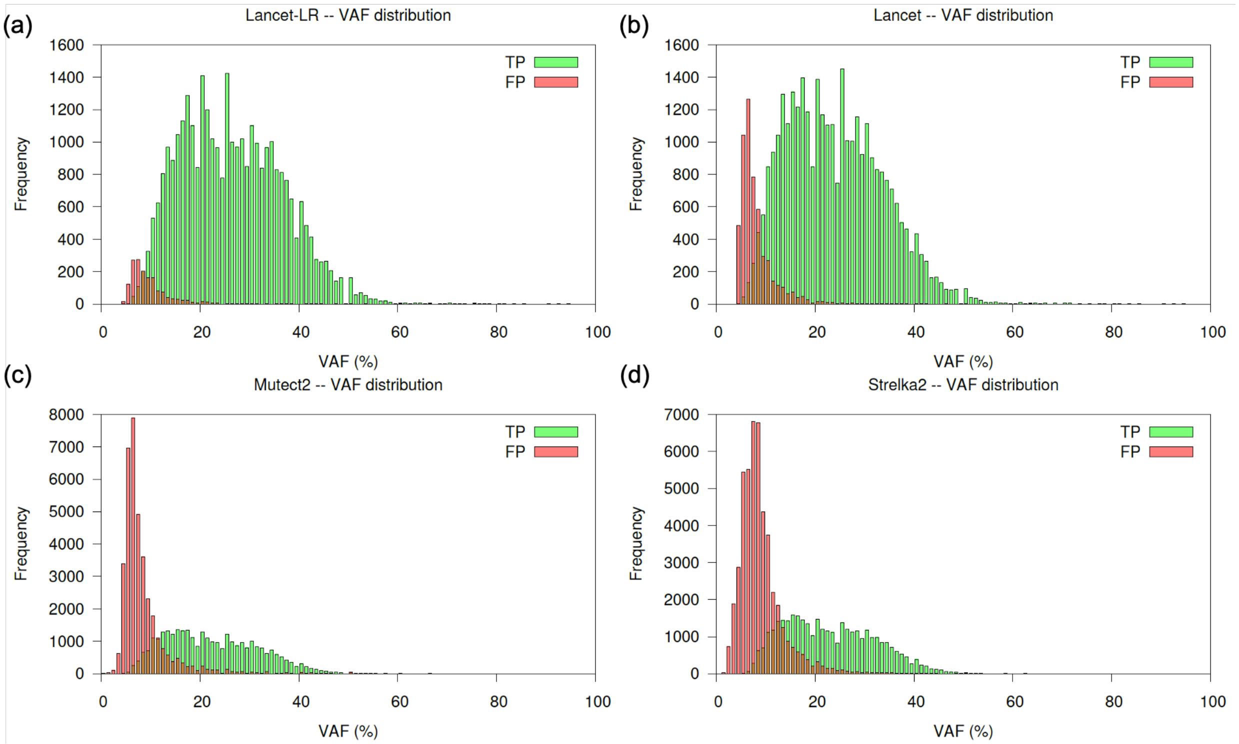


**Figure S8. Variant allele fractions distribution of SNVs.** Variant allele fractions of true-positive and false-positive SNVs reported by Lancet-LR (a), Lancet (b), Mutect2 (c), and Strelka2 (d) on the linked-reads virtual tumor.

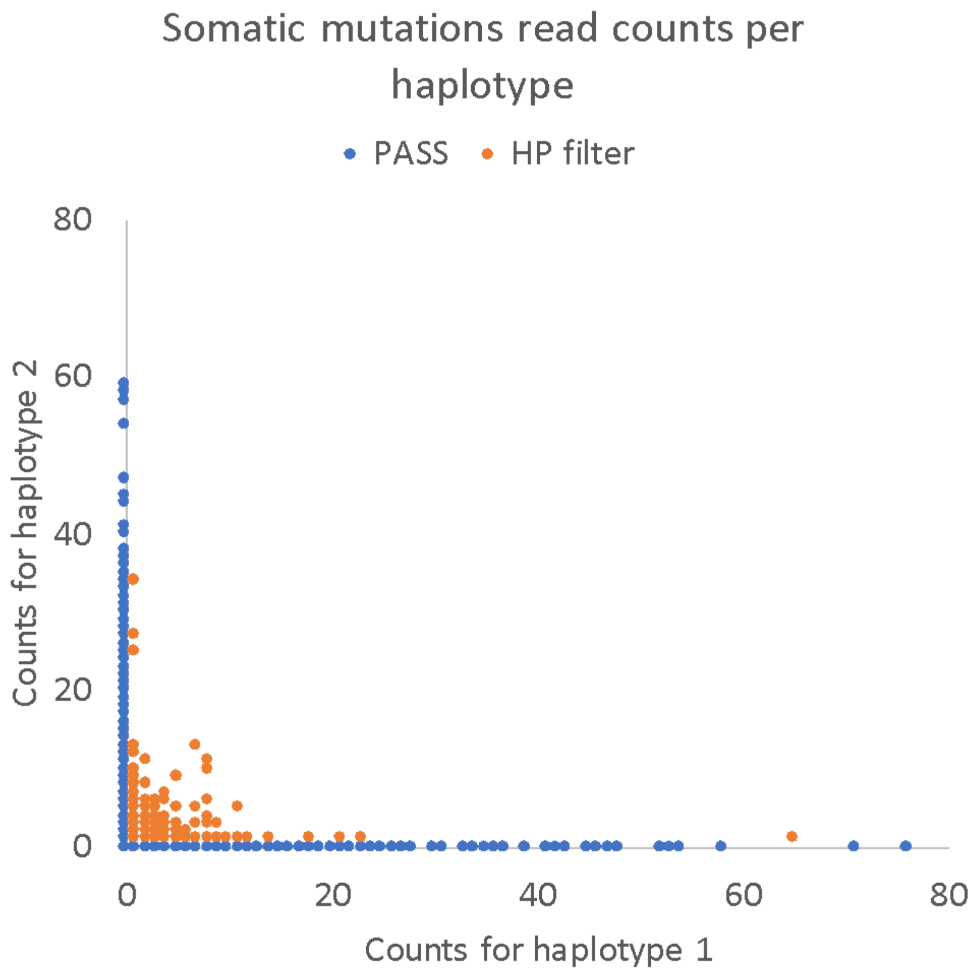


**Figure S9. Somatic mutations read counts per haplotypes.** Read counts supporting the two haplotypes show that the haplotype (HP) filter is effective in classifying low quality variants.

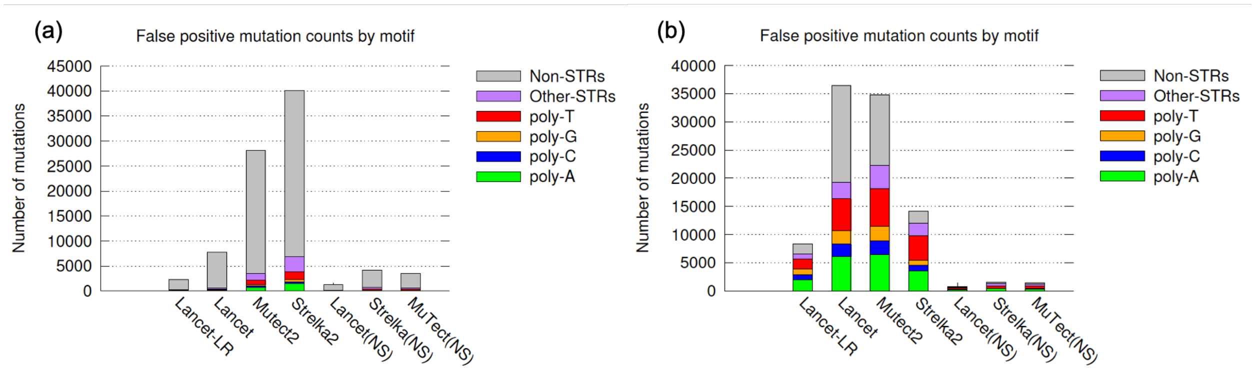


**Figure S10. False positive counts by sequence context for COLO829 cancer cell line.** Number of false positive (a) SNVs and (b) InDels reported by each caller on the COLO829 cancer cell line**.** The (NS) notation indicates results on Illumina NovaSeq sequencing data, while all other reported results are for the linked-reads data.

**Table T1.** Description of datasets used in this study.

| Dataset | Library type | Sequencing Instrument | Mean coverage | Accession/URL | Source |
|---|---|---|---|---|---|
| HCC1954 tumor | PCR free | HiSeqX | Downsampled to 30x | 6d8044f7-3f63-487c-9191-adfeed4e74d3 | TCGA mutation calling benchmark 4[1] |
| HCC1954 normal | PCR free | HiSeqX | 30x | 99d19ebd- 832c-4f2d-b97f-51743a2cf9a2 | TCGA mutation calling benchmark 4 |
| HCC1954 tumor | 10x | HiSeqX | 30x | HCC1954T_WGS_210 | 10x Genomics Datasets[2] |
| HCC1954 normal | 10x | HiSeqX | 30x | HCC1954N_WGS_210 | 10x Genomics Datasets[3] |
| COLO829 tumor | PCR free | HiSeqX | Downsampled to 30x | ERR2752450 | ENA project PRJEB27698[4] |
| COLO829 normal | PCR free | HiSeqX | 30x | ERR2752449 | ENA project PRJEB27698 |
| COLO829 tumor | 10x | NovaSeq | 30x | ERR2820167 | ENA project PRJEB27698 |
| COLO829 normal | 10x | NovaSeq | 30x | ERR2820166 | ENA project PRJEB27698 |
| 10x Virtual tumor | 10x | NovaSeq | 80x | ftp://ftp.nygenome.org/lancet/10x_virtual_tumors/bams/tumor.bam | NYGC FTP |
| 10x Virtual normal | 10x | NovaSeq | 40x | ftp://ftp.nygenome.org/lancet/10x_virtual_tumors/bams/normal.bam | NYGC FTP |

1 https://gdc.cancer.gov/resources-tcga-users/tcga-mutation-calling-benchmark-4-files
2 https://support.10xgenomics.com/genome-exome/datasets/2.1.0/HCC1954T_WGS_210
3 https://support.10xgenomics.com/genome-exome/datasets/2.1.0/HCC1954N_WGS_210
4 https://www.ebi.ac.uk/ena/data/view/PRJEB27698

**Table T2.** SNV & InDel results for COLO829 cancer cell line.

| | #Tool | # calls | TPs | FPs | FNs | Recall | PPV | FDR | F-score |
|---|---|---|---|---|---|---|---|---|---|
| SNV | Lancet (NS) | 34428 | 33330 | 1098 | 3804 | 0.90 | 0.97 | 0.07 | 0.93 |
| | Mutect2 (NS) | 38314 | 35158 | 3156 | 1976 | 0.95 | 0.92 | 0.07 | 0.93 |
| | Strelka2 (NS) | 41235 | 35726 | 5509 | 1408 | 0.96 | 0.87 | 0.09 | 0.91 |
| | Lancet-LR | 34055 | 31799 | 2256 | 5335 | 0.86 | 0.93 | 0.11 | 0.89 |
| | Lancet | 40317 | 32836 | 7481 | 4298 | 0.88 | 0.81 | 0.15 | 0.85 |
| | Mutect2 | 52568 | 34645 | 17923 | 2489 | 0.93 | 0.66 | 0.23 | 0.77 |
| | Strelka2 | 77463 | 35284 | 42179 | 1850 | 0.95 | 0.46 | 0.38 | 0.62 |
| InDel | Lancet (NS) | 1125 | 401 | 724 | 155 | 0.72 | 0.36 | 0.52 | 0.48 |
| | Strelka2 (NS) | 1985 | 502 | 1483 | 54 | 0.90 | 0.25 | 0.60 | 0.40 |
| | Mutect2 (NS) | 1876 | 475 | 1401 | 81 | 0.85 | 0.25 | 0.61 | 0.39 |
| | Lancet-LR | 8633 | 319 | 8314 | 237 | 0.57 | 0.04 | 0.93 | 0.07 |
| | Strelka2 | 14557 | 418 | 14139 | 138 | 0.75 | 0.03 | 0.94 | 0.06 |
| | Mutect2 | 33477 | 403 | 33074 | 153 | 0.72 | 0.01 | 0.98 | 0.02 |
| | Lancet | 36779 | 348 | 36431 | 208 | 0.63 | 0.01 | 0.98 | 0.02 |

F1-score: harmonic mean of precision and recall, 2 × (precision × recall)/(precision + recall).
NS: NovaSeq Illumina data. Entries sorted by the F-score metric.

**Table T3.** SNV & InDel results for HCC1954 breast cancer cell line.

| | #Tool | # calls | TPs | FPs | FNs | Recall | PPV | FDR | F-score |
|---|---|---|---|---|---|---|---|---|---|
| SNV | Lancet-LR | 16318 | 12827 | 3491 | 6635 | 0.66 | 0.78 | 0.21 | 0.72 |
| | Lancet | 19386 | 13129 | 6257 | 6333 | 0.67 | 0.68 | 0.32 | 0.67 |
| | Mutect2 | 56722 | 15845 | 40877 | 3617 | 0.81 | 0.28 | 0.72 | 0.41 |
| | Strelka2 | 68311 | 15430 | 52881 | 4032 | 0.79 | 0.22 | 0.77 | 0.35 |
| | Lancet (HiSeq) | 17668 | 15839 | 1829 | 3623 | 0.81 | 0.89 | 0.10 | 0.85 |
| InDel | Lancet-LR | 5413 | 628 | 4785 | 1093 | 0.36 | 0.11 | 0.88 | 0.17 |
| | Lancet | 16555 | 724 | 15831 | 997 | 0.42 | 0.04 | 0.95 | 0.08 |
| | Mutect2 | 31663 | 1019 | 30644 | 702 | 0.59 | 0.03 | 0.96 | 0.06 |
| | Strelka2 | 15649 | 1026 | 14623 | 695 | 0.59 | 0.06 | 0.93 | 0.12 |
| | Lancet (HiSeq) | 1452 | 923 | 529 | 798 | 0.53 | 0.63 | 0.36 | 0.58 |

F1-score: harmonic mean of precision and recall, 2 × (precision × recall)/(precision + recall).

### Section S1. Enhancements to support linked-reads in Lancet

#### Linked-reads integration in the De Bruijn graph

We have extended and augmented the Lancet colored De Bruijn graph structures to seamlessly integrate the long-range and phasing information provided by the linked-reads. Each node (*k*-mer) has been extended to store the set of barcodes (BX tag) and haplotype group (HP tag) of the linked-reads from where each *k*-mer was extracted. Specialized algorithms have been implemented to take advantage of this information across multiple key components of the method. For example, when threading through the graph, a *barcode-aware* coverage is computed by counting the number of distinct barcodes associated to the *k*-mers/nodes in the path. Similarly to the way overlapping reads in Illumina fragments are typically handled, the *barcode-aware* coverage avoids overcounting the contribution of reads sharing the same barcode molecules with uneven coverage (**Supplementary Figure S3**). Similarly, the haplotype assignments of the *k*-mer in the graph are used to flag somatic variants that are not found in conjunction with only one haplotype.

#### Command line options and usage

Lancet new linked-reads mode of operation can be easily activated using the following command line options:

- --linked-reads, -J : linked-reads analysis mode
- --primary-alignment-only, -I : only use primary alignments for variant calling

The complete list of all other available parameters and flags can be found on the Lancet GitHub page (https://github.com/nygenome/lancet). LongRanger BAMs are directly supported (https://support.10xgenomics.com/genome-exome/software/pipelines/latest/what-is-long-ranger), however, for improved accuracy, we highly recommend to pre-process the BAMs with the MarkDuplicates program from Picard Tools (https://broadinstitute.github.io/picard/), which marks PCR duplicates more accurately than LongRanger.

A typical command line to process 10x Genomics linked-reads data with Lancet looks like this:

```
lancet –J –I --tumor <BAM> --normal <BAM> --ref <FASTA> --reg <chr:start-end>
```

#### VCF format

Lancet generates the list of variants in VCF format (v4.1). All variants (SNVs and indels either shared, specific to the tumor, or specific to the normal) are exported in output. Following VCF conventions, high quality variants are flagged as PASS in the FILTER column. For non-PASS variants the FILTER info reports the list of filters that each variant does not satisfy. Specifically, in linked-reads mode, low quality variants supported by reads from multiple haplotypes are flagged with "MultiHP":

```
##FILTER=<ID=MultiHP,Description="Supporting reads from multiple haplotypes based on linked-reads analysis">
```

The output VCF also lists all the barcodes (BX) supporting the reference and alt alleles in each of the called variants. This information can be used downstream to identify somatic mutations co-occurring on the same haplotype, chromosome copy, or subclone and so phased together.

#### Section S2. Linked-Reads Virtual Tumor generation

In order to generate the "linked-reads virtual tumors", we used the strategy described in Cibulskis. et. al., 2013 (doi:10.1038/nbt.2514) with some modifications to handle the linked-reads data and the associated barcode information. Specifically, we sequenced the HapMap samples NA12891[5] and three technical replicates of NA12892[6] at ~80x and ~60x coverage respectively using the 10x Genomics libraries protocol on Illumina NovaSeq at the New York Genome Center. Each dataset was then processed with the LongRanger pipeline and Picard MarkDuplicates was run on the LongRanger output BAMs. We then spiked-in reads from NA12891 into NA12892 at sites that were known germline variants (generated by GATKv3.5 best practices pipeline) for which NA12891 was homozygous ALT (GT: 1/1) and NA12892 was homozygous REF (GT: 0/0). At each of the selected sites, a select number of reads (*N*) from NA12892 were replaced with *N* reads from NA12891, where *N* was determined by a binomial distribution using allelic fractions of 0.15 and 0.3. While replacing the reads from NA12891 to NA12892, we retained all the alignment information of the source sample (NA12891) but match the molecule and barcode information of the destination sample (NA12892). This procedure allows to keep the raw data and alignment information as realistic as possible without disrupting the molecule structure of the destination sample (**Supplementary Fig. S11**). Through this process, a number of additional variants, present in the spiked-in reads at different locations then the pre-selected ones, are unintentionally introduced; these variants are cataloged in a "to ignore" list and not used during benchmarking evaluation. Overall this procedure resulted in the spike-in of 38,062 SNVs and 3,654 InDels (**Supplementary Fig. S12**).

[5] https://www.coriell.org/0/Sections/Search/Sample_Detail.aspx?Ref=GM12891
[6] https://www.coriell.org/0/Sections/Search/Sample_Detail.aspx?Ref=GM12892

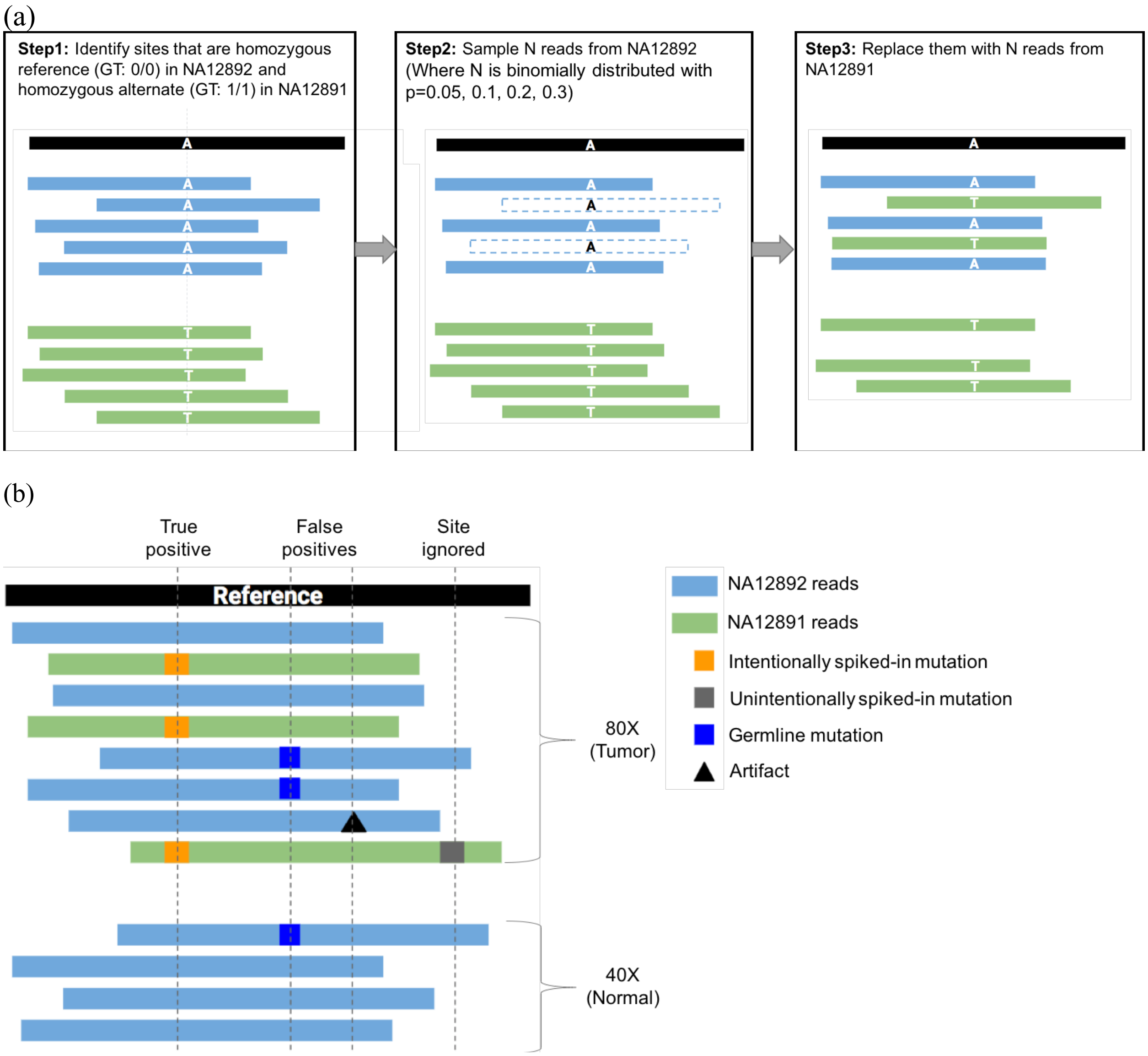


**Figure S11. Virtual tumors as benchmarking datasets for evaluation of somatic variant callers**. Strategies for (a) creation of virtual tumors and (b) evaluation of somatic variant callers run on them.

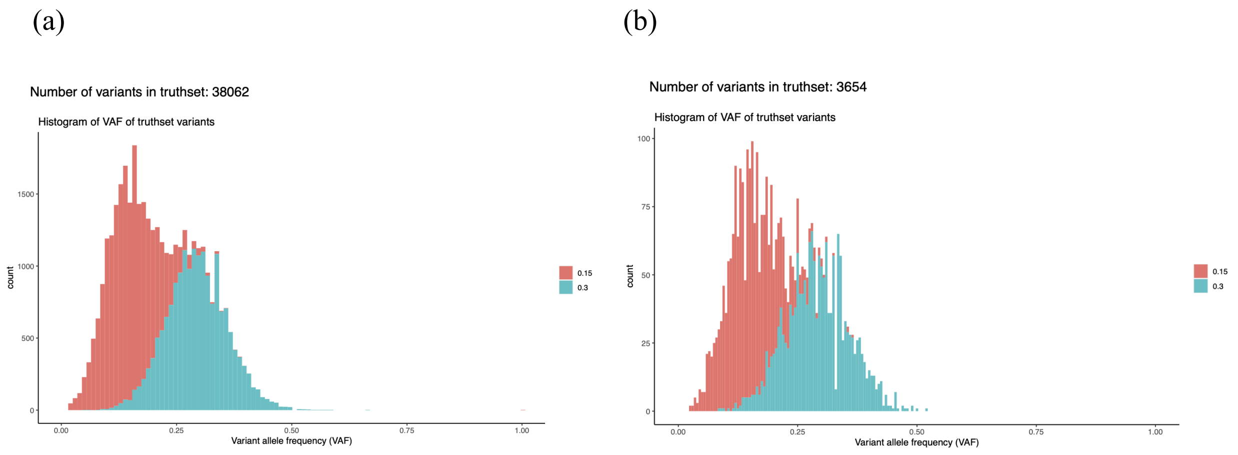


**Figure S12. Variant allele frequency (VAF) distribution of the spiked-in variants.** Histograms of VAF of spiked-in (a) SNVs (n=38,062) and (b) InDels (n=3,654) in the linked-reads virtual tumor. The colors represent the allelic fractions used as probabilities (*p*) for the binomial distribution used to spike the variants.

### Section S3. Description of the benchmarking workflow

#### Variant callers

- Mutect2 packaged along with GATK v4.0.5.1
- Strelka2 v2.9.3
- Lancet v1.1.0

#### Commands used

- NYGC Cancer WGS pipeline v6.0[7] (Arora *et al.*, 2019) was used to run Mutect2 and Strelka2.
- Lancet was run with default parameters by including the new linked read specific flags, "--linked-reads" and "--primary-alignments-only" when benchmarking 10x datasets.

#### Variant calls filtering and evaluation

For each caller we kept only the PASS somatic variants within the autosomes together with chromosomes X, Y and sorted the variant calls, from highest quality to the lowest, according to the quality score reported by each method in the final VCF file ("FisherScore" for Lancet, "SomaticEVS" for Strelka2, "TLOD" for MuTect2). Due to possible ambiguous representation of indels around microsatellites and other simple repeats, indel calls from the various callers, including the truth and ignore set variants, were left normalized before comparing them during the benchmarking process. When comparing calls to the truth set or across the different methods, we matched two variants (SNV or indels) if they shared the same genomic coordinates (chromosome and start position), as well as if they have the exact same sequences (both in size and base pair composition) in the reference and alternative alleles. Precision/recall values along the curve are computed for each tool by processing the somatic calls in the sorted order according to the specific tool's quality score.

[7] https://www.nygenome.org/bioinformatics/software/nygc-cancer-pipeline
